# Relating brain connectivity with persistent symptoms in pediatric concussion

**DOI:** 10.1101/539825

**Authors:** Kartik K. Iyer, Karen M. Barlow, Brian Brooks, Zahra Ofoghi, Andrew Zalesky, Luca Cocchi

## Abstract

Persistent post-concussion symptoms (PCS) in children following a mild traumatic brain injury (mTBI) is a growing public health concern. There is a pressing need to understand the neural underpinning of PCS. Here, we examined whole-brain functional connectivity from resting-state fMRI with behavioral assessments in a cohort of 110 children with mTBI. Children with mTBI and controls had similar levels of connectivity. PCS symptoms and behaviors including poor cognition and sleep were associated with connectivity within functional brain networks. The identification of a single “positive-negative” dimension linking connectivity with behaviors enables better prognosis and stratification towards personalized therapeutic interventions.

## INTRODUCTION

Traumatic brain injury (TBI) in children has a worrying yearly incidence of 798–1373 per 100,000^1^. Although 90% of these are mild TBIs (mTBI), children are slower to recover and 11 to 30% of children still experience persistent post-concussion symptoms (PCS) three months later^2^. Children with persistent PCS typically exhibit a heterogeneous collection of psychological, physical, and behavioral symptoms^3^. Childhood PCS is further compounded by a lack of targeted clinical interventions or management strategies that alleviates prolonged symptom presentation and disturbed brain activity. As a result, PCS may impact the neurodevelopment of a child and the long-term functional outcome^4, 5^.

Persistent PCS symptoms have been associated with alterations to structural and functional brain networks^4, 6–8^. Notably, deficits in functional connectivity within circumscribed brain networks have been linked to distinct behavioral symptoms including fatigue^4^. Despite this knowledge, the link between symptoms, behavior, and patterns of functional brain network connectivity in children with persistent PCS remains largely unknown. This information is crucial to reveal the neural underpinning of heterogeneous symptoms, develop new diagnostic and prognostic markers, as well as to implement targeted therapeutic interventions that are personalized to each patient.

In this study, we recruited a large sample of children diagnosed with persistent PCS and used a validated single multivariate analysis to assess the relationship between resting functional brain connectivity and a constellation of heterogeneous symptoms and behaviors that commonly arise from the condition^9^. Based on previous work in healthy^9^ and clinical populations^10^, we expected symptoms and behaviors to link with abnormal patterns of whole-brain connectivity along a single or multiple dimensions. Our results suggest that a single dimension links whole-brain connectivity, behavior, and symptoms of persistent PCS. The analysis of associations within this dimension provides novel key insights on neural and behavioral factors supporting persistent pediatric PCS.

## MATERIALS AND METHODS

### Subjects

One hundred and ten children with a confirmed diagnosis of concussion/mTBI were recruited together with 20 well matched healthy controls. Specifically, children diagnosed with a concussion/mTBI and assessed as having an increase in PCS symptoms compared with pre-injury status at 4-weeks post injury were included in the study. We also applied the following exclusion criteria: (i) a child with a prior significant medical history or previous concussion within 3 months; (ii) use of medication that would likely affect neuroimaging participation and/or sleep; and (iii) an inability to complete questionnaires and/or neuropsychological evaluation (e.g. non-English speaking background). Furthermore, ten children with mTBI were excluded following neuroimaging data quality control (see *Neuroimaging* section for details). Demographic and clinical characteristics are shown in Table 1. Ethical approval was granted by the University of Calgary Conjoint Health Research Ethics Board (Canada) and the University of Queensland (Australia).

**Table 1.**
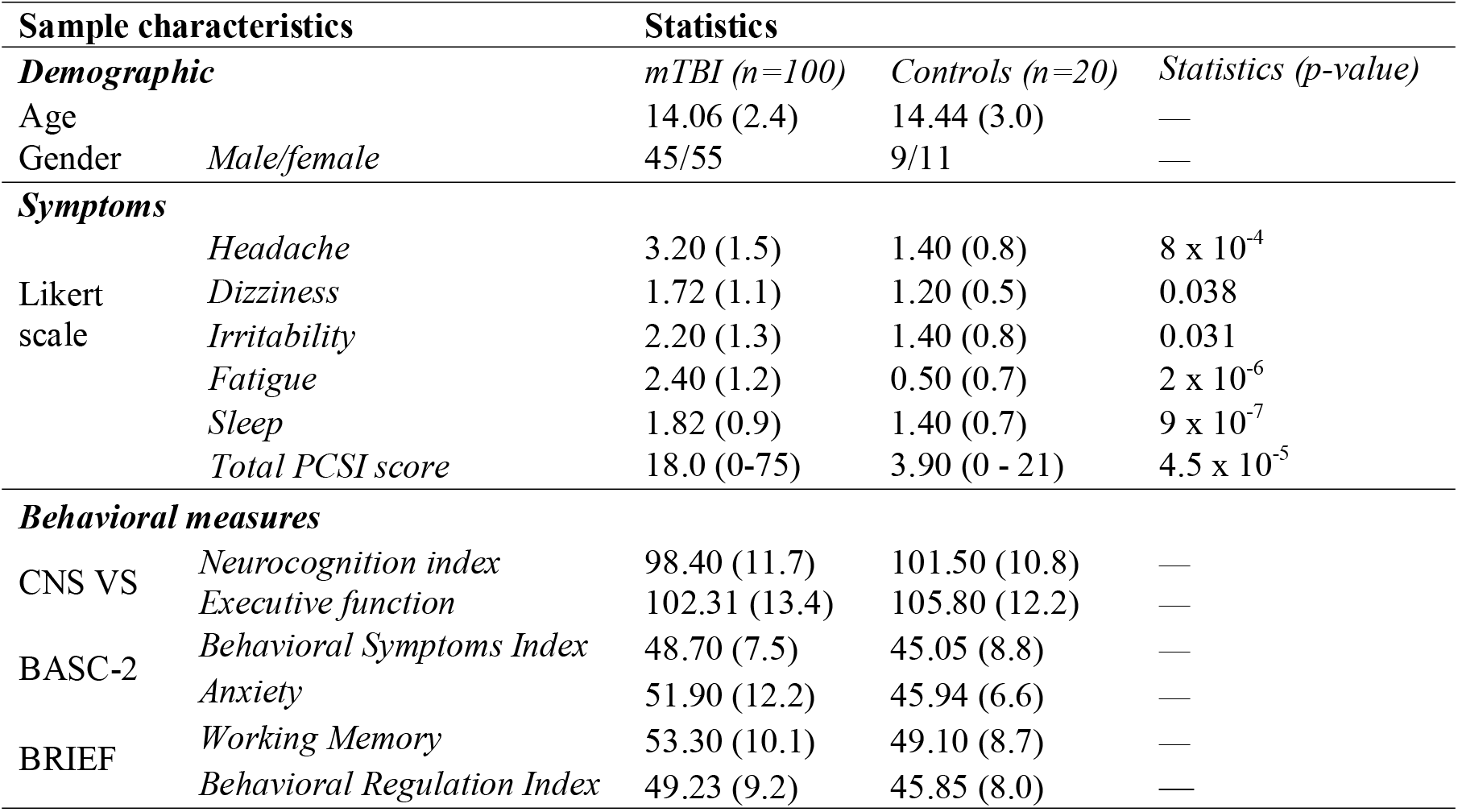
Participant demographics including sample size, age profile for the two groups and gender distribution; with mean and standard deviation (SD) where appropriate. Regarding symptoms, the median symptom scale and PCSI (Likert-scale) for the whole cohort are shown. CNS VS: Computerized Neurocognitive Software Vital Signs, Standard score; BASC-2: Behavior Assessment System for Children, second edition (T scores); BRIEF: Behavior Rating Inventory Executive Function (T Scores). The Chi Square test was used to assess putative group difference in Age whereas the Wilcoxon rank sum test was used to assess differences in symptoms and behavioral measures. “—” indicates non-significant values.

### Clinical and behavioral measures

Pediatric PCS commonly presents as a heterogeneous set of clinical and behavioral manifestations^2, 5^. However, several symptoms and behaviors are commonly associated with the condition and routinely assessed in clinical settings. To maximize the clinical relevance and translational value of our study, we focused on these common presentations of pediatric PCS. Specifically, we recorded the Post Concussion Symptom Inventory — Parent report (PCSI) which is a sensitive and standardized clinical rating scale used to summate key domains of post-concussive symptoms and behaviors such as headaches, mood and sleep disturbance^11^.

Behavioral and neuropsychological measures commonly used to monitor persistent PCS severity and recovery were also employed: (i) CNS Vital Signs Neurocognition Index: An overall performance index of cognitive and attentional response to tasks^12^; (ii) Behavior Assessment System for Children-second edition (parent report): a validated measure of a child’s adaptive and problem behaviors. This includes an anxiety subdomain measuring a child’s tendency to appear nervous, fearful, or worried^13^; and (iii) Behavior Rating Inventory of Executive Function^14^ (parent report): A well validated measure of executive function that includes measurements of working memory capacity and behavioral regulation (emotion and cognitive control).

### Neuroimaging

Functional brain connectivity during resting state was measured using a 3-T GE scanner (32-channel head coil). Gradient echo EPI data were recorded for five minutes and ten seconds (TE = 30 ms, TR = 2000 ms, flip angle = 90, FOV = 23, 150 volumes, slice thickness 3.6 mm). This relatively short acquisition time achieved an optimal trade-off between the need for sufficient data and the necessity of minimizing burden for unwell children^15^. A structural scan (T1) was also acquired using an oblique plane with 0.8 mm slice thickness (flip angle = 10°, inversion time: 600 ms, FOV = 24.0).

Pre-processing using DPARSFA V4.4 included: slice timing, realignment, segmentation via DARTEL^16^, normalization using DARTEL (3 mm^3^), smoothing using DARTEL (9 mm^3^), regression of nuisance covariates (linear trends, head motion parameters using Friston 24, cerebrospinal fluid, white matter signals), and bandpass filter between 0.01 to 0.1 Hz. Frame-wise displacements (FD) exceeding a threshold of 0.4 mm were censored^17^. Quality control analysis showed that micro-head movements (FD) were not significantly different between groups (unpaired t-test, *p* = 0.29). Subjects with less than 95% of data remaining after the removal of contaminated volumes (FD > 0.4, including one preceding and two following volumes) were excluded (*n* = 10).

A validated whole-brain parcellation comprising 214 regions was used to compute Pearson correlation in functional magnetic resonance imaging (fMRI) signals between all pairs of regions^18^. This recent brain parcellation was adopted because it has been rigorously tested with resting-state fMRI and specifies superior functional homogeneity. Resulting connectivity matrices were fisher-Z transformed. Each of the 214 regions was assigned to one of seven canonical resting-state networks^19^. Parcels were assigned to a given brain network based on a minimum Euclidian difference criterion.

For each child, in both groups, we calculated the median connectivity across all pairs of regions comprising a given network, as well as pairs of regions linking the bilateral frontoparietal networks with the default mode network^19^ (**Fig. 1**). Connectivity between the frontoparietal and default mode networks was estimated as deregulation across network communication has been suggested to play a key role in supporting pathological cognitive functions^20–22^.

**Figure 1.**
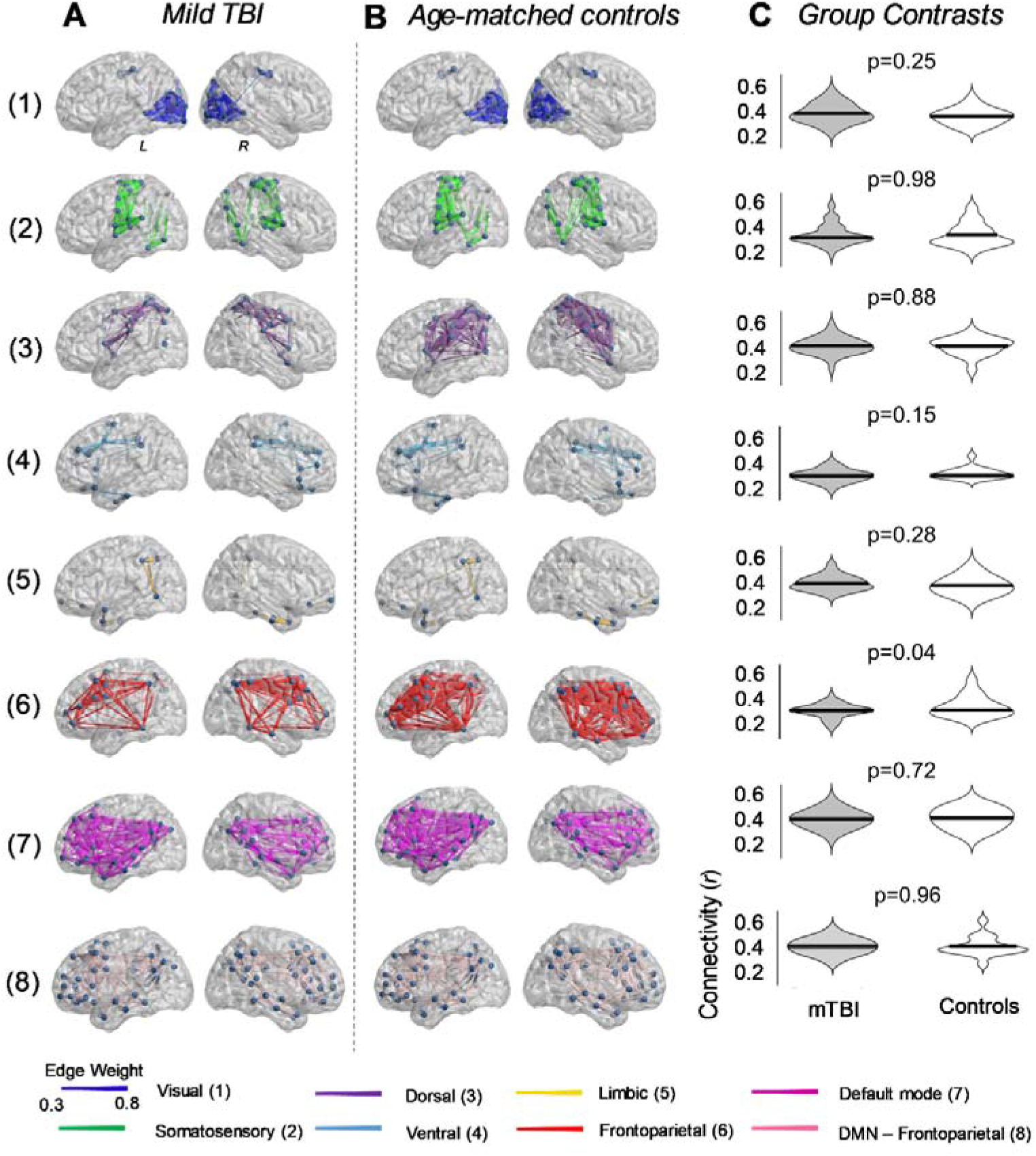
Comparison of resting-state functional connectivity in children with mild TBI. Resting-state functional connectivity was calculated in seven validated resting-state brain networks^19^, as well as between the frontoparietal and default mode networks. **(A)** Anatomical representation of the seven functional networks, segregated according to group. Patterns of connectivity between the frontoparietal and default mode networks are also depicted (row 8). Edge diameter indicates group-averaged functional connectivity strength (Pearson correlation). **(B)** Distribution of functional connectivity strength between mTBI and healthy controls. Functional connectivity did not significantly differ between the two groups (two-tailed unpaired t-test; FDR corrected).

Canonical correlation analysis (CCA) was used to independently test whether inter-individual variation in within- and between-network connectivity estimates was significantly associated with persistent PCS symptoms and behavior^9^ in mTBI and control cohorts. Permutation testing (*n* = 15,000) and bootstrapping was used to: (i) assess if a significant global relationship (mode) was greater than chance^10^ (*p*_FWE_ < 0.05, family-wise error correction, FWE), (ii) ascribe statistical significance to distinct CCA network and behavior weights. Additionally, the following checks were performed to ensure that: (i) subjects with any missing behavioral and clinical measures were excluded for dataset completeness, (ii) no non-linear relationships and outliers could affect the CCA outputs, and (iii) no multicollinearities or singularities existed amongst variables. To ensure that our findings were robust, we also performed a train-test validation analysis. Using a standard *k*-fold cross validation method, we fitted 10 CCAs on randomly selected “train” datasets comprising 70% of the subjects. The network and behavioral weights estimated from each CCA were subsequently used to perform “test” CCAs on 30% of the left-out participants. Note that the canonical weights were not refitted for the test data. Instead, that weights estimated on the training data were applied to the test data and the strength of main mode of variation was determined.

## RESULTS

Within-network functional connectivity measured in the seven brain networks did not significantly differ between groups (**Fig. 1**). Connectivity between the frontoparietal and default mode brain network was also similar between the two groups.

We next examined whether inter-individual variation in median functional connectivity strength was significantly associated with the expression of symptoms and behaviors observed in children with mTBI. Eight children were further excluded from analysis due to incomplete behavioral measurements (*n* = 92).

CCA results showed a single significant mode of population variation linking connectivity, symptoms, and behavioral measures in persistent PCS (**Fig. 2A**, *r* = 0.65, *p*_FWE_ = 0.035). Furthermore, permutation testing showed that the significant association detected by the CCA was above chance level. Additionally, the remaining CCA modes were non-significant (see **Supplementary Figure 1**). The cross validation (*k*-fold) analysis confirmed that the main CCA mode estimated from randomly selected train data sets (*n* = 65 out of 92 patients) retained the same network and behavior relationships observed in **Fig. 2A** (with weighted Pearson correlations within the range *r* = 0.65 to 0.80). The significance of the main CCA mode was confirmed in left-out datasets, by applying the CCA network and behavior weights from our trained set to smaller subsets of data (*n* = 27 out of 92 patients) and re-running CCA analysis with the same permutation testing. An independent CCA analysis for age-matched controls on the relevant behavioral and clinical measures with connectivity across networks did not reveal a significant mode of covariation (*p*_FWE_ = 0.88).

**Figure 2.**
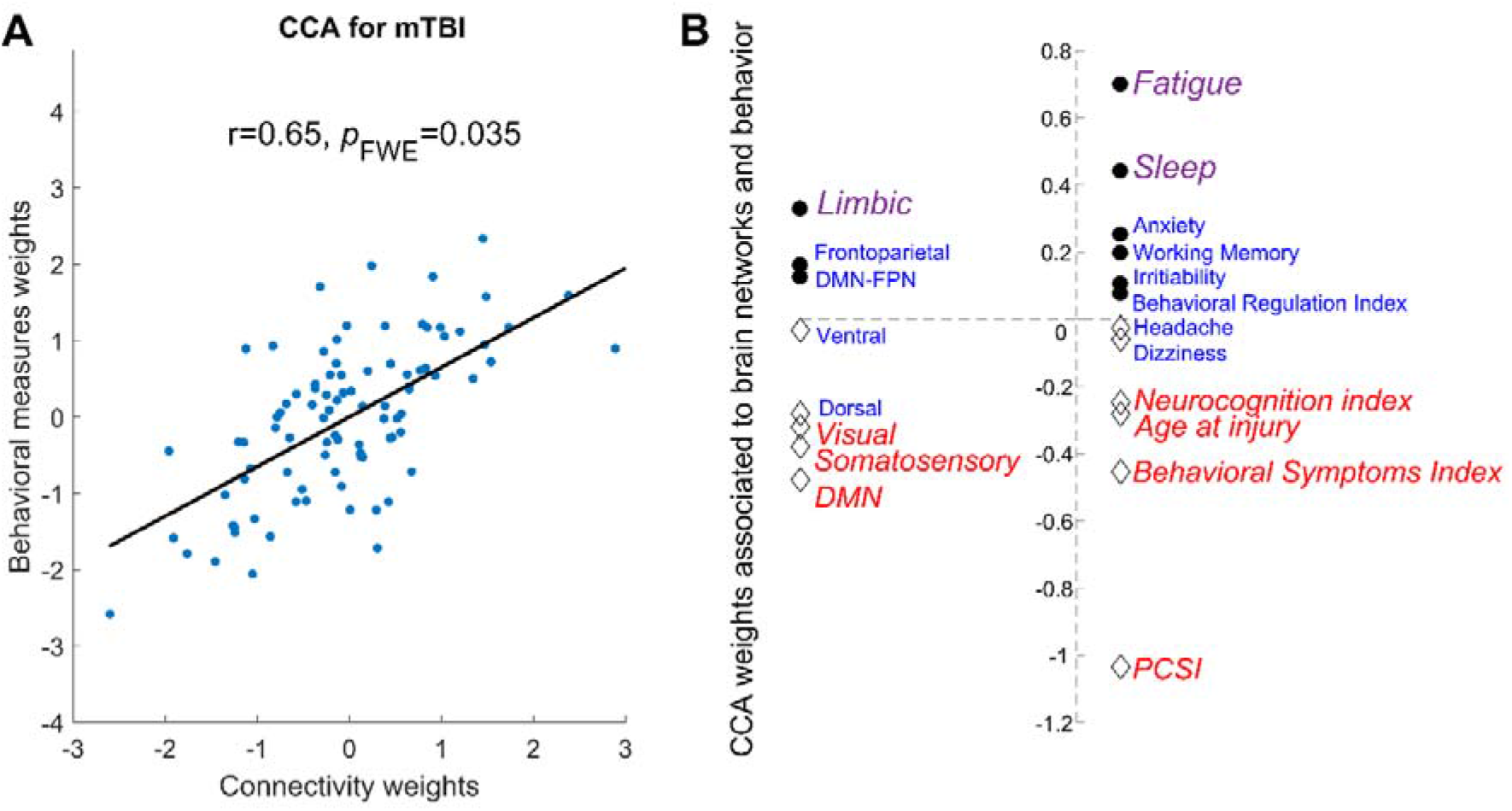
Canonical correlation analysis (CCA) assessing the multivariate association between resting-state connectivity, symptoms, and behavior in children with persistent PCS. **(A)** A single significant mode of population variation was found between resting-state connectivity in the seven brain networks (including connectivity between frontoparietal and default mode networks) and behavioral measures and symptoms of persistent PCS. **(B)** Specific contribution of different brain networks connectivity (left) and behavioral measures (right) to the CCA mode assessed using permutation testing. Significance (*p*_FWE_<0.05) loading is highlighted by *italic*, with the relative contribution of each factor indexed by both the text size (larger equates to greater contribution) and color (purple = significant positive load, blue = neutral, red = significant negative load). PCSI= Post Concussion Symptom Inventory (total score).

In mTBI, poorer neurocognition, older age at injury, worse behavioral symptoms, higher PCSI scores, and reduced connectivity in visual, somatosensory, and default mode networks (DMN) negatively loaded to the global mode (**Fig. 2B**, red colored text). In contrast, higher symptoms of fatigue, increased sleeping problems, and higher connectivity within the limbic network showed a significant positive load (**Fig. 2B**, purple colored text).

## DISCUSSION

The lack of knowledge regarding the complex association between whole-brain network connectivity and diverse clinical symptoms and behaviors is a major impediment to the development of efficient management strategies for persistent PCS in pediatric populations. To address this knowledge gap, we used multivariate statistics to link neuroimaging, clinical symptoms, and behavioral data in a large cohort of well characterized children with post-concussive symptoms following mTBI. Results showed that individual variations in resting-state functional connectivity patterns in the mTBI cohort are associated to the expression of symptoms and behavior along a single positive to negative dimension (**Fig. 2**). Decrease in connectivity within default mode, visual and somatosensory networks, as well as cognitive and emotional problems loaded negatively on the dimension. Conversely, higher connectivity within the limbic network, along with poorer sleep quality, and higher fatigue loaded positively. These results advance knowledge on the neural underpinning of symptoms and behaviors associated to pediatric PCS^4^, facilitating the planning of targeted interventions aiming to reduce the burden of mTBI following injury^23^.

### Linking whole-brain networks with symptoms and behavior defining PCS

Resting-state brain networks are thought to recapitulate a fundamental organizational principle of the brain^24^. Such networks have been identified across a range of tasks and are involved in diverse brain functions including interoception, attention, and cognitives reasoning^24–26^. The characterization of resting-state brain connectivity offers neurophysiological information to parse variability in the capability of individuals to recover from brain injuries^4, 27^. Results from our study highlighted how normal variability in median connectivity strength within major resting-state networks arising from mTBI relate to symptoms severity and behavior changes across a PCS cohort with mixed recovery trajectories. Moreover, our CCA results build on prior findings in pediatric mTBI showing that brain network activity plays a key role in mediating recovery from PCS symptoms and behaviors^4, 6, 8^.

Decreased functional connectivity between regions of the DMN has previously been associated to poorer cognition, specifically in individuals with traumatic brain injuries (TBI)^4, 7, 28^. Increased DMN connectivity in a state of rest has, on the other hand, been linked with better cognitive functions^25^. Our results support these earlier reports and the notion that DMN activity is a global marker of persistent PCS outcomes^4^. The coherent negative load between decreases in visual and somatosensory network connectivity and symptoms of cognitive and behavioral control detected here is also in line with the proposition that TBI in younger populations may have a significant adverse effect on sensory and visual systems^8^.

Our results also indicated a significant functional link between higher symptoms of fatigue, poorer sleep quality, and increased connectivity within the limbic network in pediatric mTBI. This functional link is associated to decreased cognitive functions and DMN connectivity (opposite end of the dimension, **Fig 2B**, red colored text). The relationship between sleep behavior and limbic activity is consistent with the known role of the limbic system in supporting and regulating sleep functions^29^. In mTBI, poorer sleep quality and cognitive outcomes have been associated with increases in thalamic connectivity (limbic network) and decreases to DMN connectivity^30^. Moreover, the opposing connectivity patterns in limbic and DMN connectivity observed in our study have been shown to normalize in pediatric cohorts as cognition and sleep improve^27, 29^. In addition to support and tie together previous research findings, current results provide a neurobiological validation of clinical observations suggesting a mutual benefit in restoring sleep and cognitive functions.

### Concluding remarks and further avenues

The present work provides a novel insight on a complex clinical problem by explicating the functional links between whole-brain network activity and heterogeneous symptoms of pediatric persistent PCS^2, 3^. Our study cannot exclude the possibility that the detected brain behavioral associations are present in disorders other than mTBI. However, the results support the notion that the severity of specific symptoms and behaviors defining PCS can be indexed using whole-brain connectivity measures, which may facilitate monitoring of responses to targeted neurophysiological and behavioral interventions. The interdependencies between brain network connectivity, symptoms, and behavior observed here highlight the benefits of personalizing therapeutic interventions based on a child’s PCS profile. Such targeted interventions could employ either (or a combination of): (i) non-invasive brain stimulation to selectively increase or decrease resting-state functional connectivity; (ii) targeted behavioral interventions aiming to improve sleep and reduce fatigue; and (iii) cognitive remediation to facilitate the recovery of cognitive functions. Insights gained throughout this work also offer avenues towards the identification of novel diagnostic and prognostic markers.

## SUPPLEMENTARY FIGURES

**Supplementary Figure 1.**
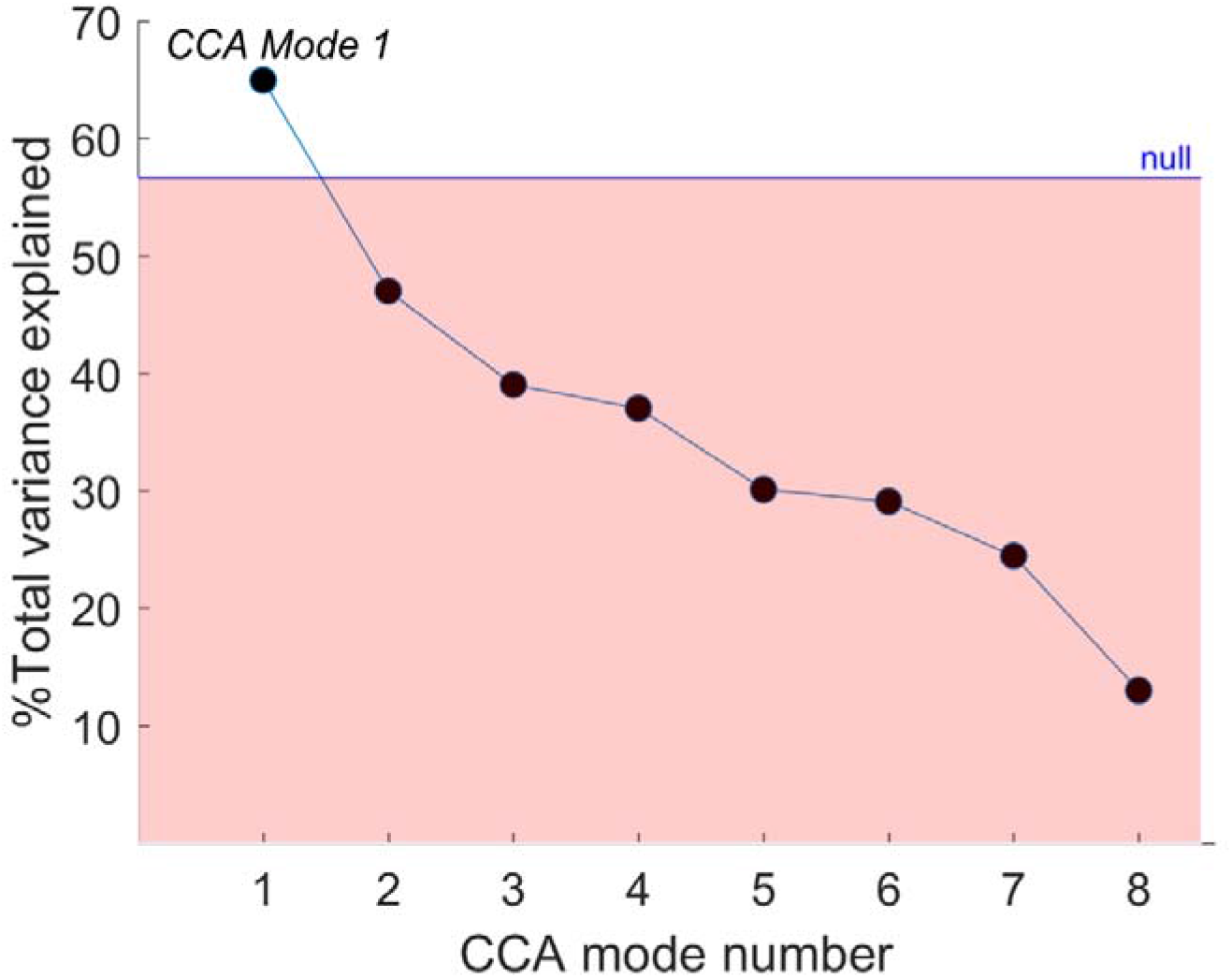
Percentage of total variance explained by our CCA analysis. We observed one significant mode of covariation (mode 1). This mode explained 65% of the total variance in the PCS cohort. The remaining seven modes did not reach statistical significance and had explained the following variance: 47%, 39%, 37%, 30%, 29%, 24%, 13%. The median total variance explained via testing with null models (15,000 permutations) was 56% (indicated by the blue line).

## ACKNOWLEDGEMENTS

K.K.I acknowledges funding support from the Brain Foundation. K.M.B. acknowledges funding support from the Canadian Institutes of Health Research (grant number: 293375). L.C. is supported by the Australian National Health Medical Research Council (L.C. 1099082 and 1138711). The authors also thank Dr Luke Hearne for feedback regarding the interpretation of the findings.

## AUTHOR CONTRIBUTIONS

K.K.I. performed all aspects of pre-processing, analysis, interpretation of results, writing of the manuscript and evaluation of findings. K.M.B. conceived, designed, acquired data for the study and provided clinical interpretation of the findings. B.B. designed and assisted with the evaluation of behavioral assessments and clinical interpretation of the findings. Z.O. was involved with acquisition and pre-processing of data. A.Z. assisted with analysis, statistical inferences and evaluation of findings. L.C. performed all aspects of pre-processing, analysis, interpretation of results, writing of the manuscript and evaluation of findings.

